# How many sirtuin genes are out there? evolution of sirtuin genes in vertebrates with a description of a new family member*

**DOI:** 10.1101/2020.07.17.209510

**Authors:** Juan C. Opazo, Michael W. Vandewege, Federico G. Hoffmann, Kattina Zavala, Catalina Meléndez, Charlotte Luchsinger, Viviana A. Cavieres, Luis Vargas-Chacoff, Francisco J. Morera, Patricia V. Burgos, Cheril Tapia-Rojas, Gonzalo A. Mardones

## Abstract

Studying the evolutionary history of gene families is a challenging and exciting task with a wide range of implications. In addition to exploring fundamental questions about the origin and evolution of genes, disentangling their evolution is also critical to those who do functional/structural studies to allow a deeper and more precise interpretation of their results in an evolutionary context. The sirtuin gene family is a group of genes that are involved in a variety of biological functions mostly related to aging. Their duplicative history is an open question, as well as the definition of the repertoire of sirtuin genes among vertebrates. Our results show a well-resolved phylogeny that represents an improvement in our understanding of the duplicative history of the sirtuin gene family. We identified a new sirtuin gene family member (*SIRT3.2*) that was apparently lost in the last common ancestor of amniotes but retained in all other groups of jawed vertebrates. According to our experimental analyses, elephant shark SIRT3.2 protein is located in mitochondria, the overexpression of which leads to an increase in cellular levels of ATP. Moreover, *in vitro* analysis demonstrated it has deacetylase activity being modulated in a similar way to mammalian SIRT3. Our results indicate that there are at least eight sirtuin paralogs among vertebrates and that all of them can be traced back to the last common ancestor of the group that existed between 676 and 615 millions of years ago.

## Introduction

The availability of whole-genome sequences in species of all main groups of vertebrates represents an opportunity to unravel the evolution of gene families. The number of genomes and their phylogenetic representativeness in the vertebrate tree of life allows performing robust inferences regarding how gene family members are related to each other and their modes of evolution (Nei and Rooney 2005). The available genomes also open an opportunity to discover new gene lineages that are not currently described maybe because they are not present in model species and/or to the absence of appropriate evolutionary analyses (Wichmann et al. 2016; Céspedes et al. 2017; Himmel et al. 2020). Gene copy number variation could be seen as a natural experiment (Albertson et al. 2009) that could help understand the evolutionary fate of duplicated genes, as individuals with different repertoires can, in principle, fulfill the biological functions with a different combination of paralogs (Gitelman 2007).

The sirtuin gene family, class III of histone deacetylase enzymes (HDACs), is a group of genes that in deuterostomes (the group that includes chordates, echinoderms, and hemichordates), is composed of seven paralogs (*SIRT1-7*) grouped into four classes (Fig. 1)(Frye 2000; Frye 2006). Sirtuin genes are involved in a variety of biological functions mostly related to aging, metabolic regulation, stress response, cell cycle among others (Fig. 1) (Michan and Sinclair 2007; Greiss and Gartner 2009; Haigis and Sinclair 2010; Zhao et al. 2019; Zhang et al. 2020). All sirtuin genes have a conserved catalytic domain and variable carboxy- and amino terminal domains. Sirtuins commonly possess NAD-dependent acyl-lysine deacylase activity (most commonly deacetylase activity), while some sirtuins may have, in addition to deacetylase activity, other enzymatic activities like ADP-ribosyltransferase, desuccinylase and demalonylase (Fig. 1). Further, they are located in different subcellular compartments and associated to different biological functions (Fig. 1).

**Figure 1.**
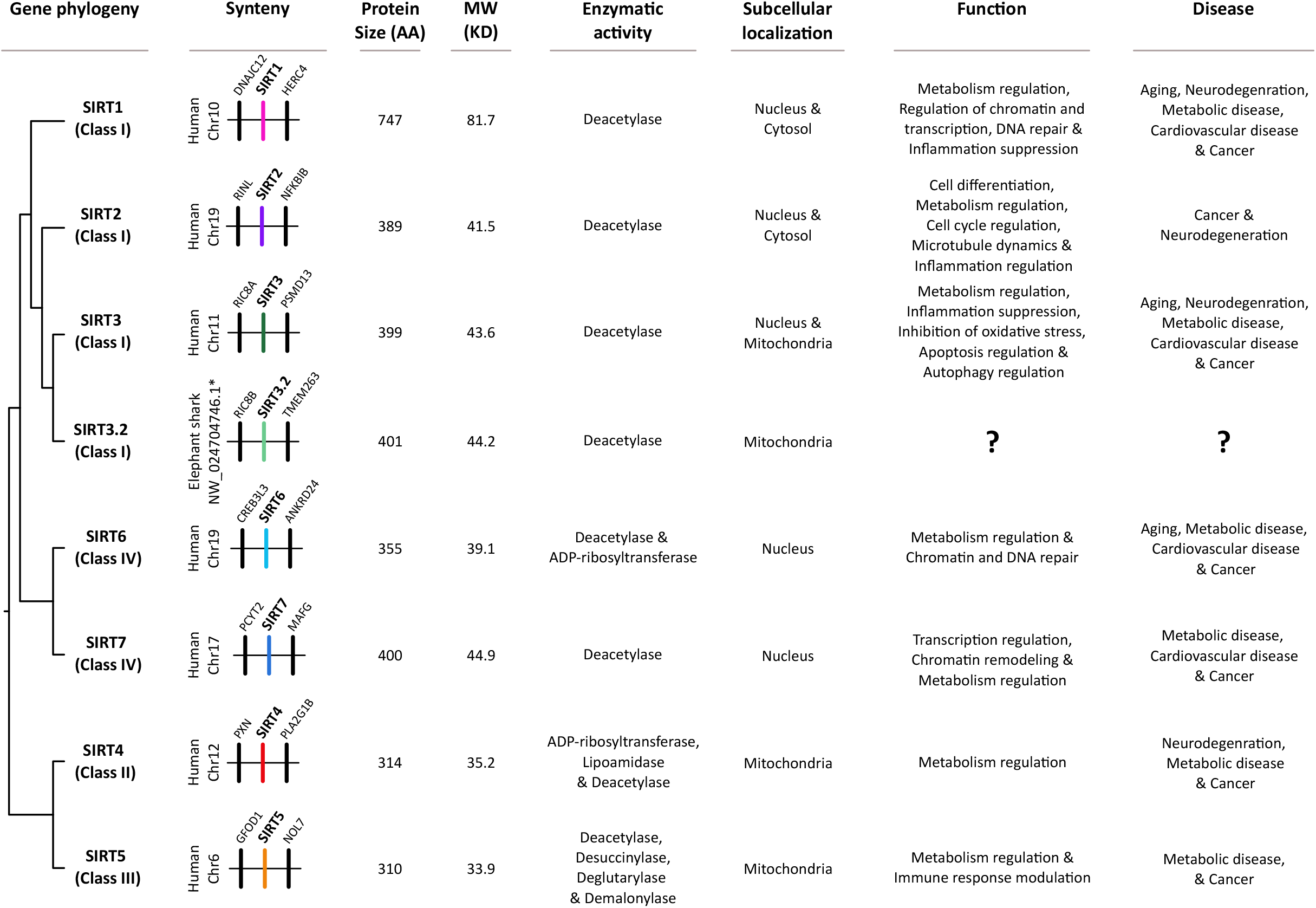
Gene phylogeny, synteny, protein size, molecular weight, enzymatic activity, subcellular localization, functions and diseases associated with sirtuin genes. Information regarding the sister group relationships among sirtuin genes was obtained from this study, synteny from ENSEMBL v.106 (Howe et al. 2021), protein size from Fujita and Yamashita (2018)(Fujita and Yamashita 2018), molecular weight from Vassilopoulos et al. (2011)(Vassilopoulos et al. 2011) while enzymatic activity, subcellular localization, functions and diseases from Zhang et al. (2020)(Zhang et al. 2020). In the case of the SIRT3.2 of the elephant shark, synteny information was obtained from the National Center for Biotechnology Information (NCBI) (Sharma et al. 2019), protein size, molecular weight, enzymatic activity and subcellular localization from this study.

The evolution of the sirtuin genes is an active area of investigation. There are multiple phylogenetic hypotheses describing evolutionary relationships among the orthologs and paralogs in the sirtuin gene family (Frye 2000; North and Verdin 2004; Frye 2006; Greiss and Gartner 2009; Pereira et al. 2011; Slade et al. 2011; Vassilopoulos et al. 2011; Costantini et al. 2013; Scholte et al. 2017; Simó-Mirabet et al. 2017; Yang et al. 2017; Rajabi et al. 2018; Kabiljo et al. 2019; Zhao et al. 2019; Gold & Sinclair 2022). Differences in the taxonomic sampling, number of orthologs and/or paralogs included, and the inclusion of outgroups are likely contributors to this variation. Additionally, studies that are focused on resolving evolutionary relationships are scarce, and in fact, most phylogenetic analyses for sirtuin genes are part of studies where the sirtuin duplicative history is a secondary goal. Additionally, and probably for similar reasons, there are no systematic efforts to characterize the full complement of sirtuin genes among vertebrates. Thus, unraveling the evolutionary history of sirtuin genes represents a challenging and exciting task with a wide range of implications. Because of their role in the aging process, these genes are of great interest. In addition to exploring fundamental questions about the origin and evolution of sirtuin genes, disentangling their evolution is also critical to understanding the diversification of functional and structural phenotypes present in the sirtuin gene family.

This study aims to take advantage of the genomic data available in public databases to advance our understanding of the diversity of vertebrate sirtuin genes and to infer its duplicative history. We also aim to characterize the subcellular localization, enzymatic activity, and mitochondrial activity of the *SIRT3.2* gene (a SIRT3-paralog gene) in the elephant shark (*Callorhinchus milii*) as a representative species. Our phylogenetic tree is in general well resolved, recovering vertebrate sirtuin genes into three clades: 1) SIRT4, SIRT5, 2) SIRT6 and SIRT7, and 3) SIRT1, SIRT2, SIRT3, and SIRT3.2. The sirtuin family member SIRT3.2 is found in jawed fish and amphibians, but is absent in earlier branching deuterostomes (e.g. cephalochordates, urochordates, echinoderms and hemichordates), is absent in cyclostomes (jawless fish), and is absent in amniotes (the group that includes mammals, birds, and reptiles). Based on how sirtuin genes are related to each other and the information which is already known for the other sirtuin family members, particularly SIRT3, we can predict that SIRT3.2 belongs to the class I, has a deacetylase activity, and is mainly located in mitochondria. Our experimental analyses confirmed our evolutionary guided inferences.

## Results and Discussion

### Vertebrate sirtuin paralogs are recovered into three main clades

We recovered the monophyly of all sirtuin family members with strong support (Fig. 2). The diversity of sirtuin genes was arranged into three main clades (Fig. 2). The first clade contains the SIRT4 and SIRT5 paralogs, the second clade contains the SIRT6 and SIRT7 paralogs, and the third clade includes the SIRT1, SIRT2, SIRT3 and SIRT3.2 gene lineages (Fig. 2). A diversity of phylogenetic arrangements for sirtuin genes have been proposed in the past (Frye 2000; Frye 2006; Greiss and Gartner 2009; Slade et al. 2011; Vassilopoulos et al. 2011; Costantini et al. 2013; Scholte et al. 2017; Simó-Mirabet et al. 2017; Yang et al. 2017; Rajabi et al. 2018; Kabiljo et al. 2019; Zhao et al. 2019; Gold & Sinclair 2022), and our results largely support the relationships proposed by Frye (2006).

**Figure 2.**
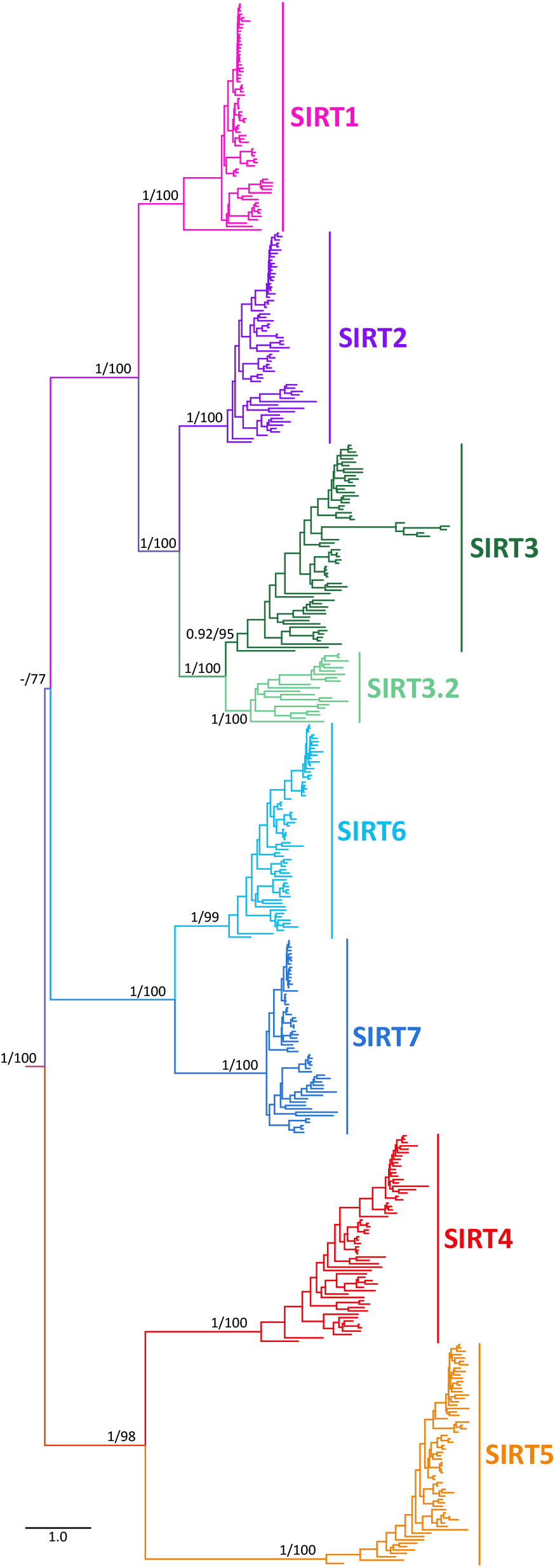
Maximum likelihood tree showing sister group relationships among sirtuin genes of vertebrates. Numbers on the nodes correspond to support from the aBayes and ultrafast bootstrap values. Nicotinamide Nucleotide Transhydrogenase (NNT) sequences from the human (*Homo sapiens*) mouse (*Mus musculus*), spotted gar (*Lepisosteus oculatus*) and zebrafish (*Danio rerio*) were used as outgroups (not shown). The scale denotes substitutions per site and colors represent vertebrate sirtuin lineages.

In the first clade we recovered the sister group relationships between SIRT4 and SIRT5 with strong support (Fig. 2). Evolutionary relationships between these two family members are still a matter of debate, as a variety of phylogenetic positions have been suggested for these paralogs (Slade et al. 2011; Vassilopoulos et al. 2011; Costantini et al. 2013; Yang et al. 2017; Rajabi et al. 2018; Gold & Sinclair 2022), and only in a fraction of the studies their sister group relationship is supported (Frye 2000; North and Verdin 2004; Frye 2006; Greiss and Gartner 2009; Hirschey 2011; Scholte et al. 2017; Simó-Mirabet et al. 2017; Kabiljo et al. 2019; Zhao et al. 2019). In the second clade, we recovered SIRT6 sister to the clade containing SIRT7 sequences with strong support (Fig. 2). The sister group relationship between SIRT6 and SIRT7 has been recovered in all examined studies (Frye 2000; North and Verdin 2004; Greiss and Gartner 2009; Hirschey 2011; Slade et al. 2011; Vassilopoulos et al. 2011; Costantini et al. 2013; Scholte et al. 2017; Simó-Mirabet et al. 2017; Yang et al. 2017; Rajabi et al. 2018; Kabiljo et al. 2019; Zhao et al. 2019; Gold & Sinclair 2022), suggesting there is robust support for the sister relationship between these sirtuin family members. In the third clade, there is a broad consensus in the literature that SIRT2 shares a common ancestor more recently in time with SIRT3 than with any other sirtuin paralog, and that the clade containing SIRT1 sequences is sister to the SIRT2/SIRT3 clade (Frye 2000; North and Verdin 2004; Frye 2006; Greiss and Gartner 2009; Hirschey 2011; Slade et al. 2011; Vassilopoulos et al. 2011; Costantini et al. 2013; Scholte et al. 2017; Simó-Mirabet et al. 2017; Yang et al. 2017; Rajabi et al. 2018; Zhao et al. 2019). We recovered the same evolutionary relationships with strong support (Fig. 2). The sister-group relationship between the SIRT6/SIRT7 and SIRT2/SIRT3/SIRT3.2/SIRT1 received moderate support (Fig. 2). The sister-group relationships among the three main sirtuin clades is something that has been difficult to resolve (Simó-Mirabet et al. 2017; Zhao et al. 2019), as it appears divergences were close in time, as evidence by the short length of the corresponding branch (Fig. 2).

In summary, we present a phylogenetic analysis based on a taxonomic sampling that included representative species from all main groups of vertebrates for all sirtuin family members. Our phylogenetic tree is in general well resolved (Fig. 2), representing an advance in our understanding of the duplicative history of the sirtuin gene family. In comparison to the phylogenetic trees currently available in the literature only one study shows the same topology as our study (Frye 2006).

#### Identification of a new sirtuin gene family member, *SIRT3.2*

In our analyses, we identified a new sirtuin family member (Fig. 2 and 3), *SIRT3.2*, which is present in a fraction of the vertebrate tree of life (Fig. 4) that was recovered sister to the *SIRT3* clade with strong support (Fig. 2 and 3). Synteny conservation provides further support to the monophyly of the *SIRT3.2* gene lineage in gnathostomes (Fig. 5a), as genes found at the 5’ side (*POLR3B, RFX4* and *RIC8B*) and 3’ side (*TMEM263, MTERF2*, and *CRY1*) of the *SIRT3.2* gene are well conserved (Fig. 5a). Synteny is also conserved in species in which the *SIRT3.2* gene was lost (Fig. 5a). Among vertebrates, we found orthologs of the *SIRT3.2* gene in representative species of cartilaginous fish, bony fish, coelacanth, lungfish and amphibians (Fig. 4). A comparison of the genomic region of the tropical clawed frog (*Xenopus tropicalis*), which possesses the *SIRT3.2* gene, with the corresponding region in the human (*Homo sapiens*), opossum (*Monodelphis domestica*), chicken (*Gallus gallus*), gharial (*Gavialis gangeticus*), red-eared slider (*Trachemys scripta*) and green anole (*Anolis carolinensis*) strongly suggests that the *SIRT3.2* gene is not present in the vertebrate lineages that these species represent (Fig. 5b). Interestingly, traces of the SIRT3.2 gene are present in the red-eared slider (*Trachemys scripta*) genome (Fig. 5b). The absence of the *SIRT3.2* gene in mammals, birds and reptiles indicates that it was probably lost in the common ancestor of the group, between 352 and 312 millions of years ago (Kumar et al. 2017). In the case of cyclostomes, we found in the sea lamprey (*Petromyzon marinus*) a chromosomal region (Chr3) that possess *TMEM263* and *POLR3B* separated by 44 base pairs, suggesting that all the genes that should be in between (*RFX4, RIC8B, SIRT3.2*) were lost in cyclostomes. Alternatively, it is possible that *SIRT3.2*, and all the other genes are present in cyclostome genomes, but not in the current genome assembly. In the past, Pereira et al. (2011), with a limited taxonomic sampling, identified a clade sister to a group containing *SIRT3* sequences. This gene lineage included sequences from three species with an evolutionary history of whole-genome duplications, two teleost fish (zebrafish, *Danio rerio*, and green spotted puffer, *Tetraodon nigroviridis*), and one amphibian (African clawed frog, *Xenopus laevis*), complicating the definition of their duplicative history. Thus, our results, including a balanced taxonomic sampling of all main vertebrate groups and appropriate phylogenetic searches, define the existence of the *SIRT3.2* gene lineage and its phyletic distribution. In agreement with Gold and Sinclair (2022), more extensive sampling in deuterostomes, now including in addition to vertebrates, urochordates, cephalochordates, echinoderms, and hemichordates, confirms that the duplication event that gave rise to SIRT3 and SIRT3.2 occurred in the last common ancestor of vertebrates (Supplementary figures 1 and 2).

**Figure 3.**
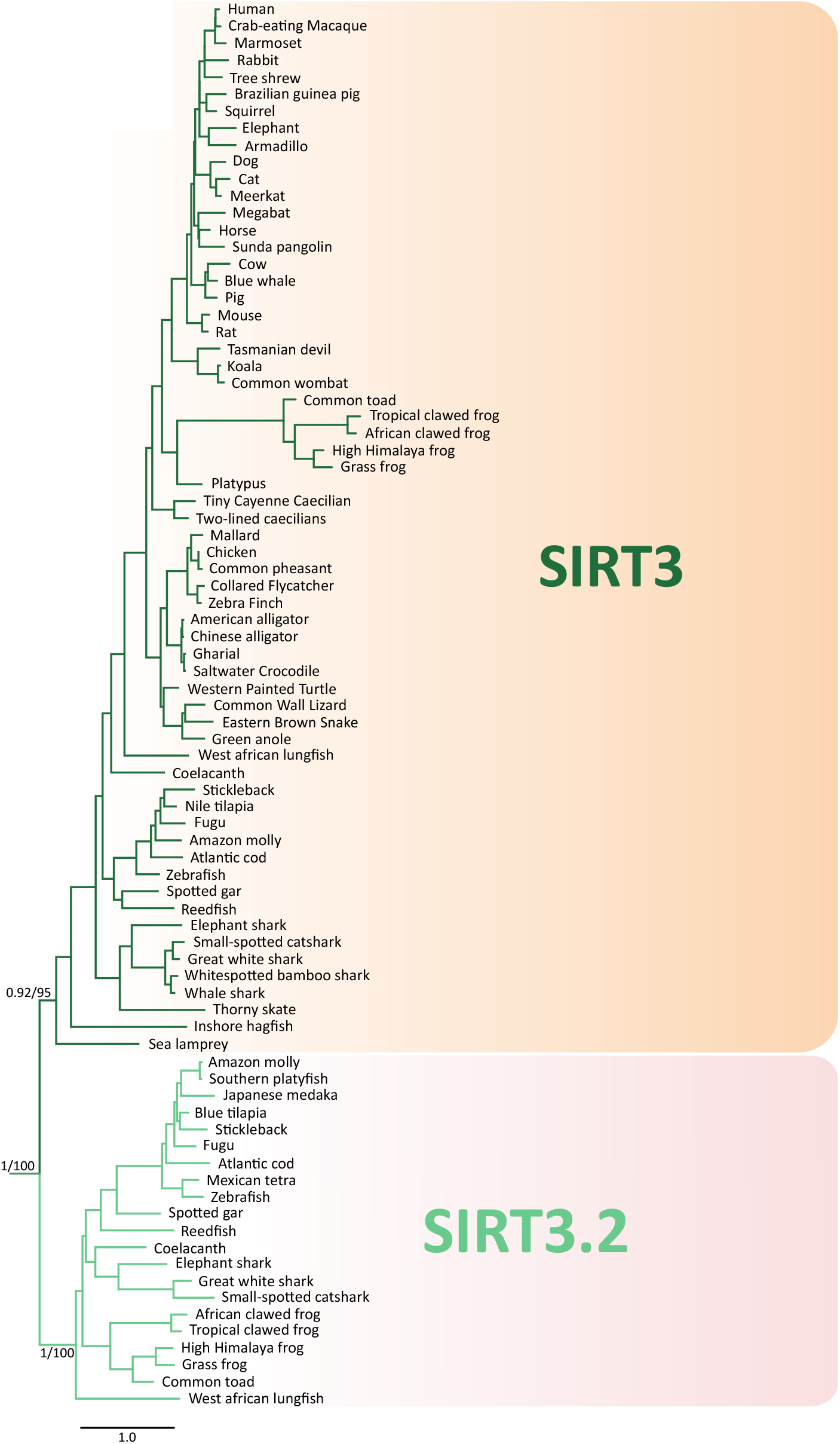
Maximum likelihood tree showing sister group relationships among SIRT3 and SIRT3.2 genes in vertebrates. Numbers on the nodes correspond to support from the aBayes and ultrafast bootstrap values. The scale denotes substitutions per site and colors represent gene lineages. This tree does not represent a novel phylogenetic analysis, it is the SIRT3/SIRT3.2 clade that was recovered from figure 2.

**Figure 4.**
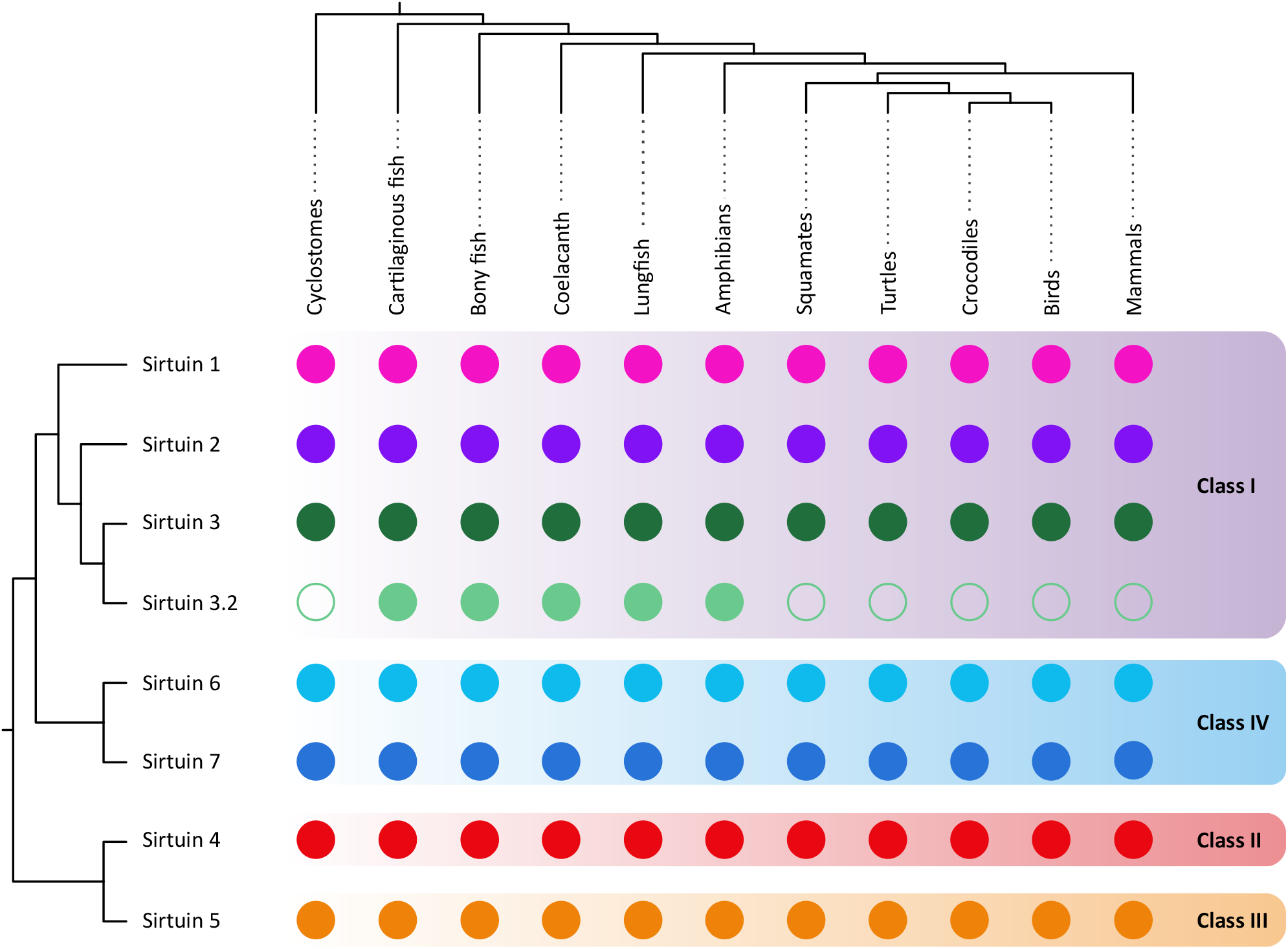
Phyletic distribution of sirtuin genes in vertebrates. The sister group relationships among sirtuin genes was obtained from this study, whereas, organismal phylogeny was obtained from the most updated phylogenetic hypotheses available in the literature (Iwabe et al. 2005; Delsuc et al. 2006; Delsuc et al. 2008; Hara et al. 2018). The colors of the circles are those used in figure 2 to define sirtuin gene lineages, whereas the classes to which the different sirtuins belong are shown with shading.

**Figure 5.**
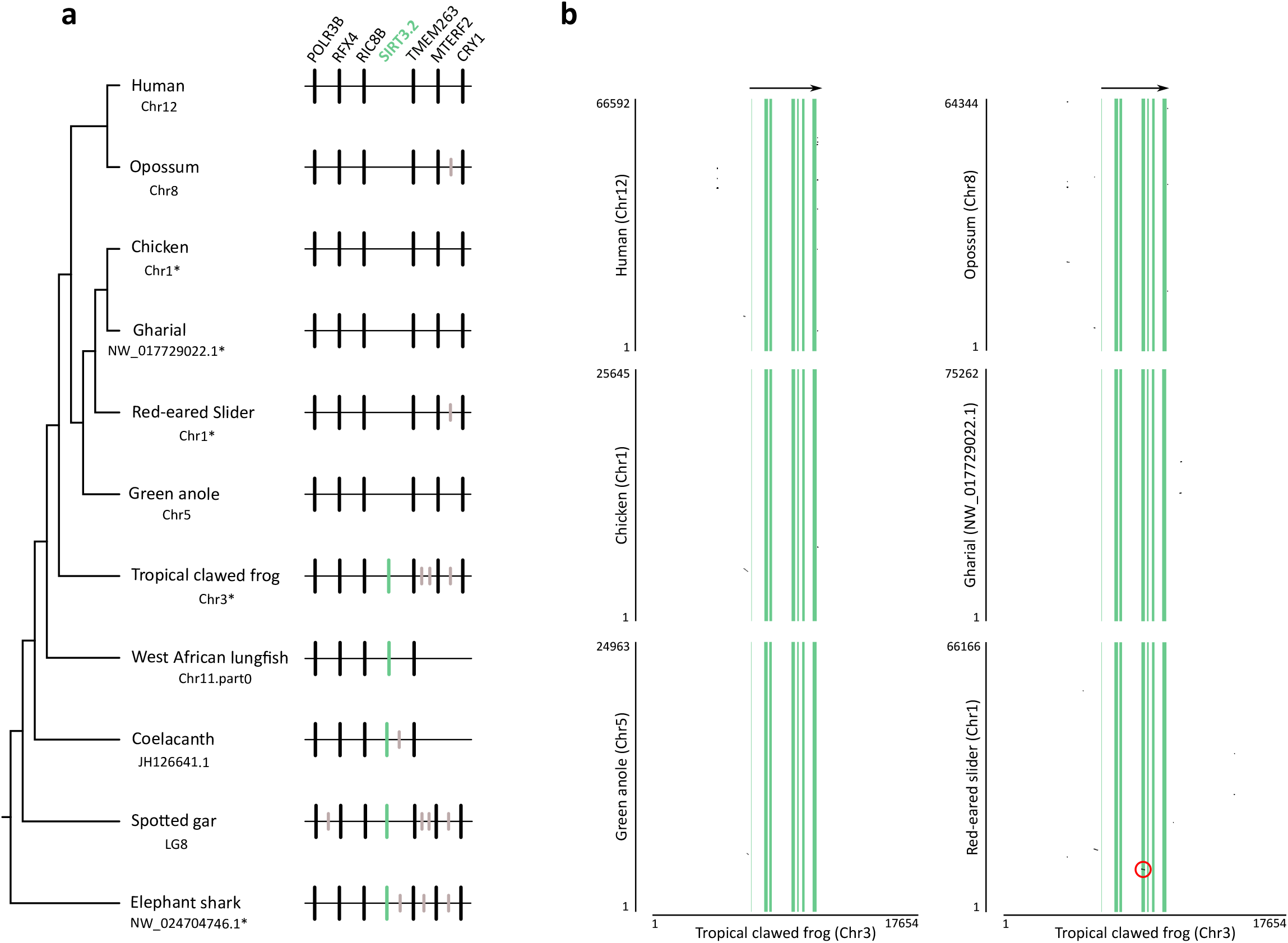
a Patterns of conserved synteny in the chromosomal region that harbor SIRT3.2 genes in gnathostomes. Asterisks denote that the orientation of the genomic piece is from 3’ to 5’, gray lines represent intervening genes that do not contribute to conserved synteny. b Pairwise dot-plot comparison of the genomic region of the tropical clawed frog (*Xenopus tropicalis*) with the corresponding region in the human (Homo sapiens), opossum (*Monodelphis domestica*), chicken (*Gallus gallus*), gharial (*Gavialis gangeticus*), red-eared slider (*Trachemys scripta*) and green anole (*Anolis carolinensis*). Light green vertical lines denote exons and regions in between are introns. Dot plots were based on the complete coding region in addition to 6.7 kb of upstream and downstream flanking sequence. The red circle highlights the vestiges of the fourth exon in the red-eared slider (*Trachemys scripta).The* genome assembly version of the species depicted in this figure is the following: Human: GRCh38.p13; Opossum: ASM229v1; Chicken: GRCg6a; Gharial: GavGan_comp1; red-eared slider: CAS_Tse_1.0; Green anole: AnoCar2.0v2; Tropical clawed frog: UCB_Xtro_10.0; Coelacanth: LatCha1; Spotted gar: LepOcu1 and Elephant shark: Callorhinchus_milii-6.1.3.

The lack of the *SIRT3.2* gene can be interpreted in different ways. We can think that the loss of *SIRT3.2* could have no physiological impact if the functional role of the lost gene is assumed by other family members, consistent with the idea that gene families possess functional redundancy (Ohno 1985; Félix and Barkoulas 2015; Albalat and Cañestro 2016). Alternatively, we can also think of gene loss as a source of adaptive evolution (Olson 1999; Nery et al. 2014; Albalat and Cañestro 2016; Helsen et al. 2020). The description of new gene lineages is not uncommon in the literature (Wichmann et al. 2016; Himmel et al. 2020; Opazo et al. 2021; Gold & Sinclair 2022), and it could be mainly attributed to the presence in non-model species and/or the absence of proper evolutionary analyses. The existence of species with different gene repertoires is an opportunity to better understand the evolutionary fate of duplicated genes (Lynch and Conery 2000) and the biological functions associated with a group of genes.

An amino acid alignment of the SIRT3.2 and SIRT3 sequences shows that the catalytic domain of SIRT3.2 is well conserved, where six amino acid positions were identified as diagnostic characters (Fig. 6). The amino acid divergence of the SIRT3.2 catalytic domain varies from 32.2% (spotted gar vs coelacanth) to 33.6% (spotted gar vs elephant shark). The same comparisons for the SIRT3 catalytic domain show slightly lower amino acid divergence values, between 24.8% (spotted gar vs coelacanth) and 31.0% (coelacanth vs elephant shark). As expected, the interparalog distance (SIRT3 vs SIRT3.2) of the catalytic domains shows higher divergence values, ranging from 40.7% (spotted gar SIRT3 vs spotted gar SIRT3.2) and 43.6% (elephant shark SIRT3 vs elephant shark SIRT3.2). Although nothing is known about the protein encoded by the SIRT3.2 gene, based on how sirtuin genes are evolutionarily related and the information already known for the other family members (Fig. 1), we can speculate that SIRT3.2 belongs to the class I, is mainly located on the mitochondria/nucleus, and has deacetylase activity. From a functional perspective it should perform functions similar to what has been described for SIRT3 (Fig. 1) (Brown et al. 2013; McDonnell et al. 2015; Zhang et al. 2020). It is important to highlight that the inferences regarding a newly discovered gene are better performed if they are phylogenetically guided.

**Figure 6.**
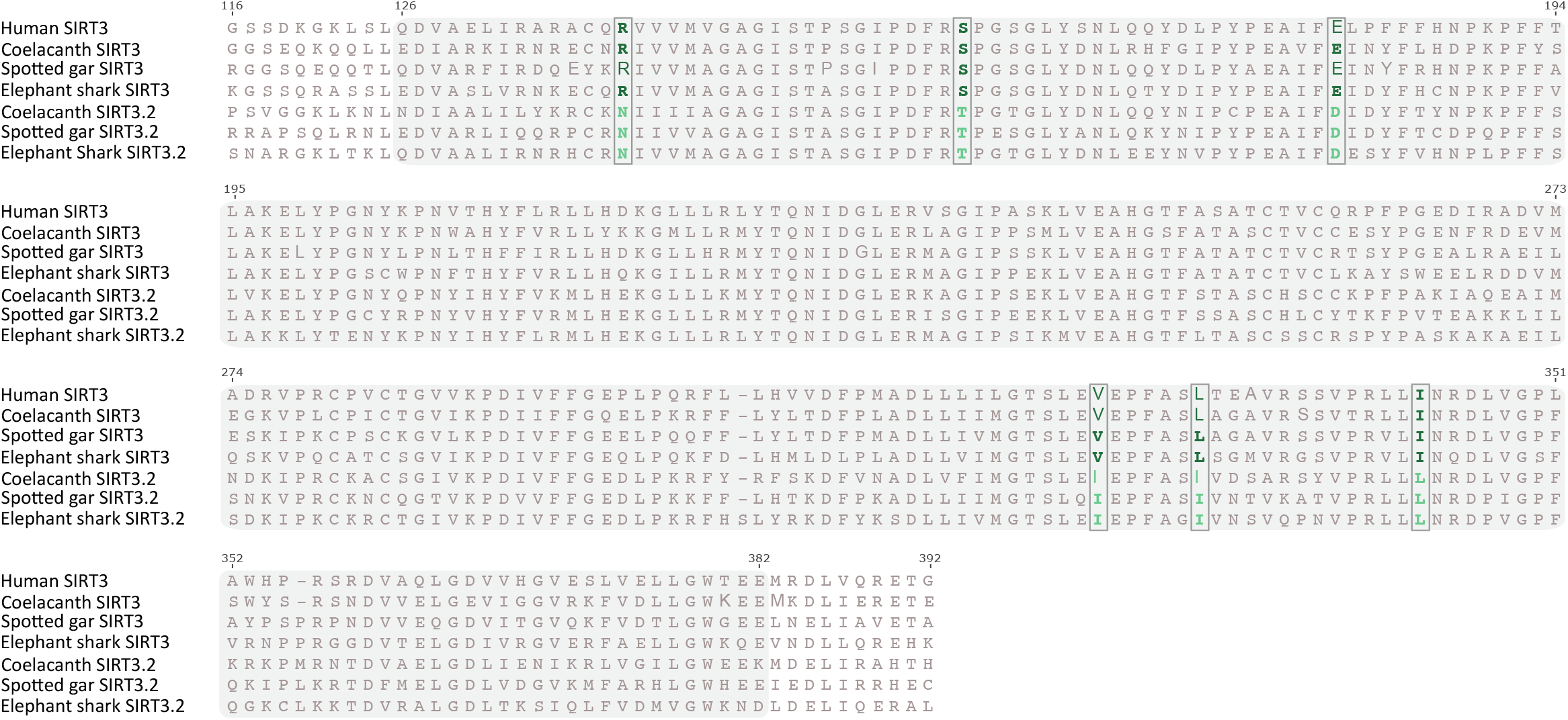
Alignment of the catalytic domain of SIRT3 in humans (*Homo sapiens*), and SIRT3 and SIRT3.2 of spotted gar (*Lepisosteus oculatus*), coelacanth (*Latimeria chalumnae*) and elephant shark (*Callorhinchus milii*). The shaded region denotes the catalytic domain. Diagnostic characters -i.e. amino acids positions that distinguish between SIRT3 and SIRT3.2 gene lineages - are indicated with a rectangle and green colors.

#### Expression pattern of sirtuin gene family members

Our next step in characterizing the *SIRT3.2* gene was to investigate whether it is transcribed and if it was, to characterize the transcription pattern. To do this, we mapped RNASeq reads to reference gene sequences of representative species of vertebrates and examined transcript abundance. In agreement with the transcription pattern reported for sirtuin genes (Stelzer et al. 2016; Kabiljo et al. 2019), our results show that although they exhibited wide variance in tissue expression, sirtuin genes are expressed in almost all tissues, including the novel *SIRT3.2* gene lineage (Fig. 7). In the case of the tropical clawed frog, the *SIRT3.2* gene is transcribed at a similar level in all tissues, other than the ovary where high transcription levels were estimated (Fig. 7). Coincident with the pattern observed in the tropical clawed frog, the elephant shark *SIRT3.2* gene is mostly transcribed in the ovary (Fig. 7). These results should be taken with caution given the lack of biological replicates, at least within species. In the case of the zebrafish, although our transcription level estimations did not follow the trend previously mentioned, an extensive study examining the expression of all sirtuin genes in zebrafish showed that the *SIRT3.2* gene in this species is mainly expressed in the ovary (Pereira et al. 2011). Thus, in addition to showing that the *SIRT3.2* gene is transcribed, our analyses allow us to suggest that this gene could be involved in biological processes associated with reproduction. This observation agrees with the literature as for SIRT3 there are many studies in which its functions related to reproduction have been reported (Zhao et al. 2016; Tatone et al. 2018; Vazquez et al. 2020; Di Emidio et al. 2021).

**Figure 7.**
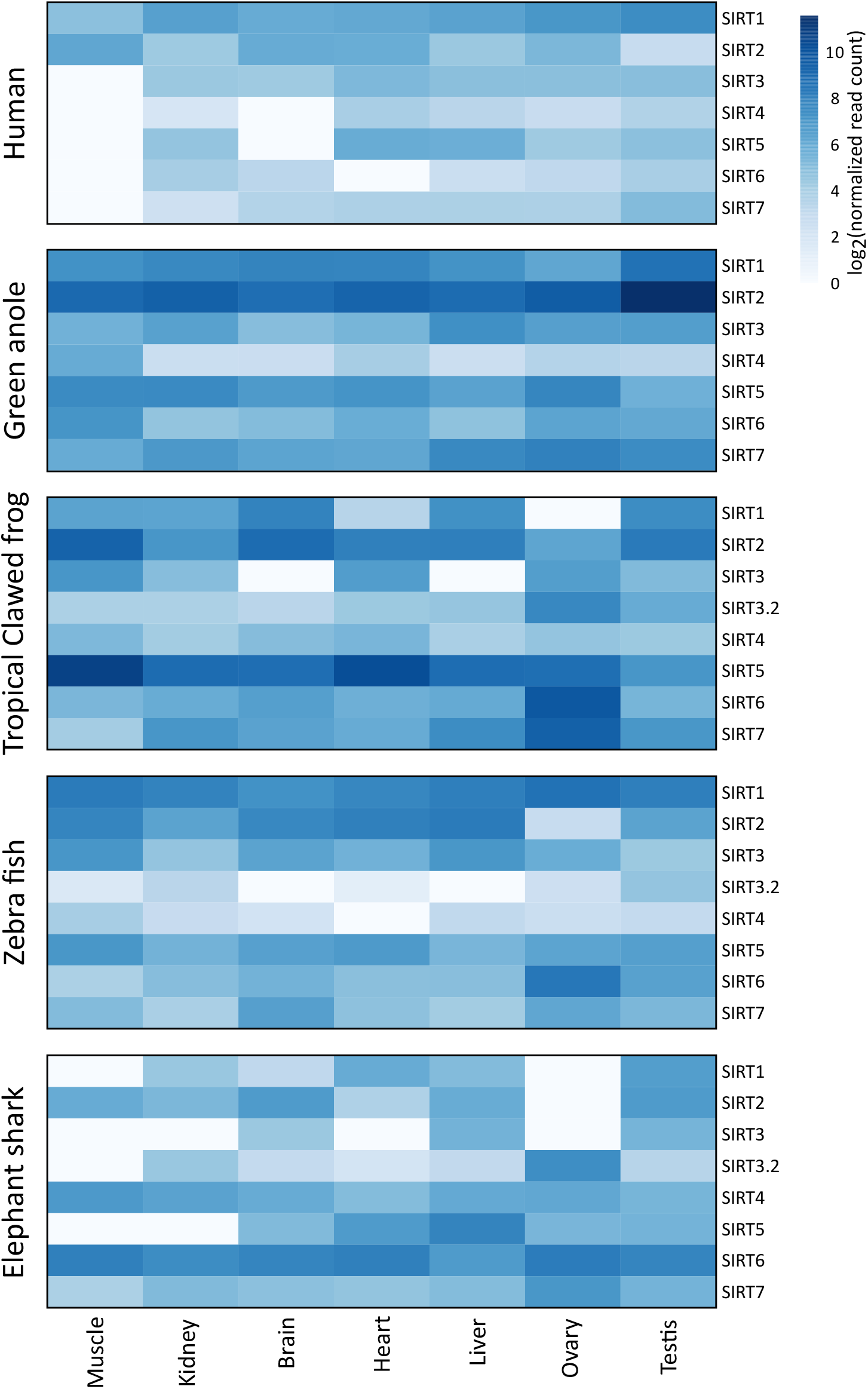
Heatmap representation of within-species relative transcriptional levels of sirtuin paralogs between 7 chosen tissues. Transcription values were calculated independently for each species and normalized over all tissues.

#### Elephant shark SIRT3.2 protein localizes to mitochondria

To further explore the structural relationship between SIRT3 and SIRT3.2 proteins, we performed an unbiased protein structure modeling of SIRT3.2 proteins from elephant shark (*Callorhinchus milii*), spotted gar (*Lepisosteus oculatus*), salmon (*Salmo salar*) and coelacanth (*Latimeria chalumnae*). Models of SIRT3.2 proteins were generated by the SWISS-MODEL workspace that used as template crystal structure models of human SIRT3: 4bvh.1.A for elephant shark, 5z94.1.A for spotted gar, 5d7n.3.A for salmon and 5bwo.1.B for coelacanth (data not shown). All models generated comprise only the central portion of the SIRT3.2 proteins where the catalytic and its regulatory regions are expected to be: amino acids G95 to Q365 for elephant shark, L115 to V388 for spotted gar, P114 to G387 for salmon, and K113 to P386 for coelacanth. The overall root mean square deviation for 272 superimposable Cα coordinates for the models of human SIRT3 (pdb: 4bvh.1.A; amino acids G121 to G392) and elephant shark SIRT3.2 (amino acids G95 to Q365) is 0.101 Å (Fig. 8a), suggesting extensive structural match. Some SIRT proteins, such as SIRT1, SIRT2 and SIRT3, possess evolutionarily conserved, distinct cellular localizations and functions, as evidenced by their analysis in several model species that include Drosophila melanogaster, Mus musculus and Homo sapiens (McBurney et al. 2003; Blander and Guarente 2004; North and Verdin 2004; Michishita et al. 2005; Haigis et al. 2006) (Fig. 1). Some sirtuins localize in more than one compartment, like SIRT1 and SIRT2 that localize to the nucleus and cytosol, and SIRT3 that localizes mainly to the mitochondrial matrix, but with a small fraction localizing to the nucleus (Scher et al. 2007). In contrast, SIRT4 and SIRT5 localize only to mitochondria, and SIRT6 and SIRT7 localize only to the nucleus. The sequence and structural comparison between SIRT3.2 and SIRT3 proteins, as well as subcellular localization prediction by LocTree3 (Goldberg et al. 2014), suggest that the localization of SIRT3.2 proteins is also in the mitochondria. To determine the subcellular localization of SIRT3.2 proteins, we chose to analyze SIRT3.2 from elephant shark. To this end, we expressed in human cultured cells full-length elephant shark SIRT3.2 tagged with three copies of the c-Myc epitope at the C-terminus (SIRT3.2-3myc). To obtain biochemical evidence of the expression of this protein, we performed immunoblot analysis with monoclonal antibody 9E10 against the c-Myc epitope, which showed the detection of an expected 48.3 kDa protein in all cell lines analyzed (Fig. 8b). Fluorescence microscopy analysis of human H4 neuroglioma cells expressing SIRT3.2-3myc showed decoration by the anti-c-Myc antibody of structures highly reminiscent of mitochondria, in addition to a faint fluorescent signal in the rest of the cytoplasm (Fig. 8c, d). Mitochondria localization of SIRT3.2-3myc was confirmed by co-localization with the fluorescent signal of MitoTracker Orange (Pearson’s correlation coefficient [r] = 0.94 ± 0.02; *n* = 20; Fig. 8c and Supplementary Fig. 3a). Importantly, SIRT3.2-3myc showed no localization in any other major compartment, as shown by lack of localization in the nucleus (detected with the nuclear stain DAPI; r = 0.21 ± 0.03; *n* = 20; Fig. 8c, d and Supplementary Fig. 3b), the Golgi apparatus (detected with antibody against Trans-Golgi network integral membrane protein 2 (TGN46) (Ponnambalam et al. 1996) (r = 0.34 ± 0.09; *n* = 20; Fig. 8c, d and Supplementary Fig. 3c), or the endoplasmic reticulum (detected with antibody against Cytoskeleton-associated protein 4 (P63) (Schweizer et al. 1993) (r = 0.47 ± 0.15; *n* = 20; Fig. 8d and Supplementary Fig. 3d). The same pattern of expression of SIRT3.2-3myc was obtained using other human cell lines, such as HeLa or MDA-MB-231 cells (data not shown). Altogether, these results indicate that SIRT3.2 proteins are targeted to the mitochondria.

**Figure 8.**
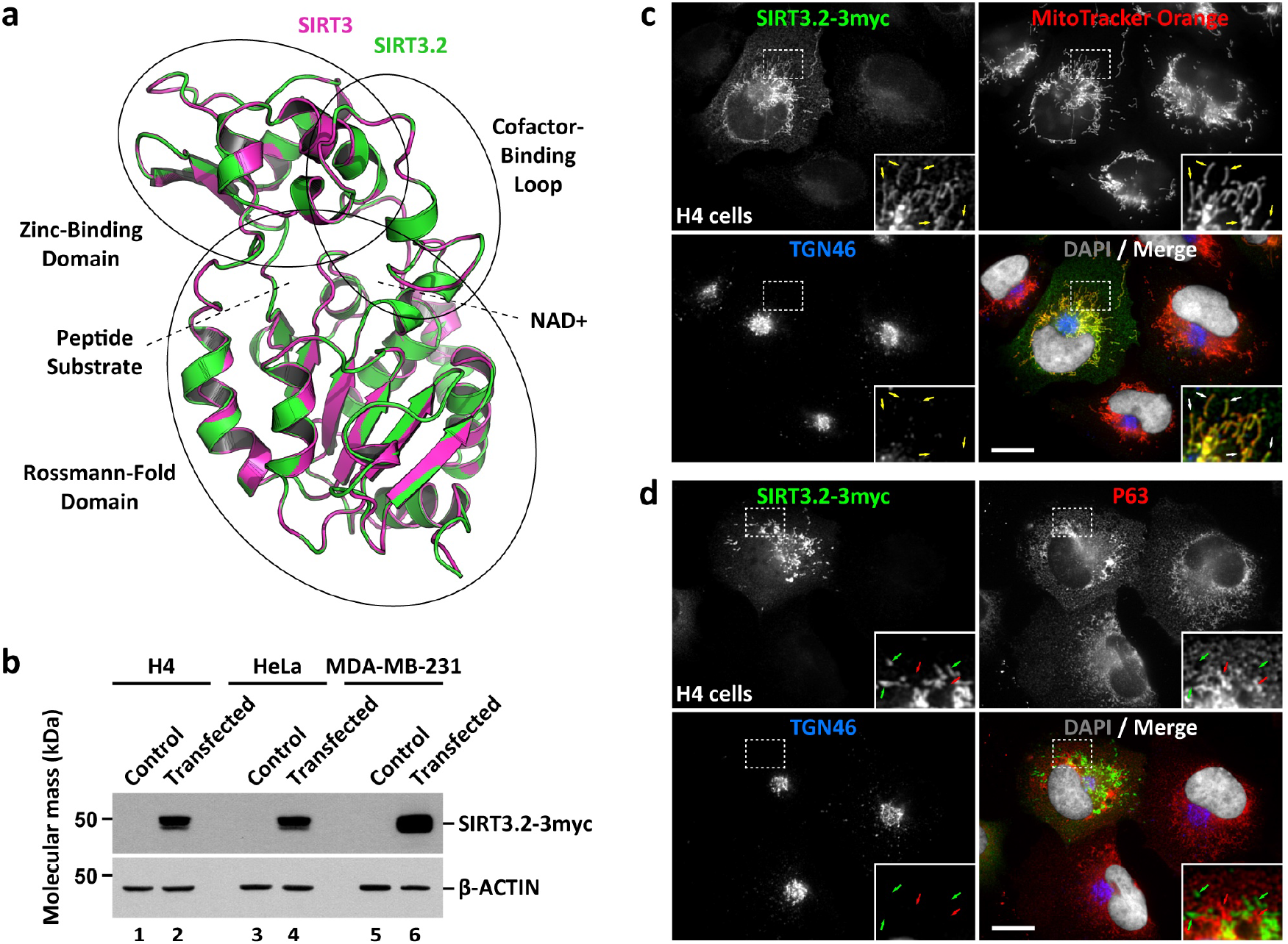
Structural model of elephant shark SIRT3.2 and expression of SIRT3.2-like-3myc in mammalian cells. a Superposition of the ribbon representations of elephant shark SIRT3.2 (green model; amino acids G95 to Q365) and human SIRT3 (magenta model; pdb: 4bvh.1.A; amino acids G121 to G392). Depicted in ellipses are the indicated functional/structural domains of SIRT3 and in dotted lines the binding sites for peptide substrates and NAD+. b The indicated cells were left untreated (Control, lanes 1, 3 and 5) or transfected to express SIRT3.2-3myc (lanes 2, 4 and 6), followed by immunoblot analysis with antibody against the c-Myc epitope or antibody against β-ACTIN used as loading control. The position of molecular mass markers is indicated on the left. SIRT3.2-3myc exhibits an electrophoretic mobility corresponding to a protein of 48.3 kDa, which is expected considering the extra amino acids of the three copies of the c-Myc epitope at the C-terminus. c and d H4 cells grown in glass coverslips were transfected to express SIRT3.2-3myc (green channel) and incubated with the mitochondrial probe MitoTracker Orange (red channel in c) or mock incubated (d). Cells were fixed, permeabilized, and double- (c) or triple-labeled (d) with mouse monoclonal antibody against the c-Myc epitope (c and d), rabbit polyclonal antibody against the endoplasmic reticulum protein Cytoskeleton-associated protein 4 (P63; d) and sheep polyclonal antibody against the Golgi apparatus protein Trans-Golgi network integral membrane protein 2 (TGN46; c and d). Secondary antibodies were Alexa-Fluor-488-conjugated donkey anti-mouse IgG (green channel), Alexa-Fluor-594-conjugated donkey anti-rabbit IgG (red channel), and Alexa-Fluor-647-conjugated donkey anti-sheep IgG (blue channel), and nuclei were stained with the DNA probe DAPI (white channel). Stained cells were examined by fluorescence microscopy. Insets: X3 magnification with yellow arrows indicating colocalization (c) and green and red arrows indicating lack of colocalization (d). Bar, 10 μm.

#### Elephant shark SIRT3.2 protein has deacetylase activity

Reversible lysine acetylation is one of the most common post-translational modifications on proteins that regulate a variety of physiological processes including gene expression, enzymatic activity, protein-protein interactions, and subcellular localization (Glozak et al. 2005). Sirtuin proteins share a catalytic core domain (North and Verdin 2004) that have NAD^+^-dependent deacetylase activity (Shahgaldi and Kahmini 2021). After translocation into the mitochondrial matrix, human SIRT3 is inactive, but after proteolytic processing of its N-terminus signal peptide it becomes active as deacetylase (Onyango et al. 2002; Schwer et al. 2002). Similar to human SIRT3, secondary structure prediction of representative SIRT3.2 proteins indicates that the first ~100 N-terminal amino acids are in a disordered domain (data not shown). Therefore, to determine if SIRT3.2 proteins also exhibit deacetylase activity, we produced in bacteria N-terminal truncated elephant shark SIRT3.2 protein (amino acids 95-372; ΔNT-SIRT3.2). To analyze the enzymatic activity of purified ΔNT-SIRT3.2, we used a commercial fluorometric deacetylation activity assay that contains an acetylated substrate peptide for recombinant human SIRT3 (Abcam). An initial analysis of activity at 37°C showed that ΔNT-SIRT3.2 possesses a deacetylase activity that is similar to that of human SIRT3, being also NAD^+^-dependent (Fig. 9a, e, f). Because the enzymatic activity of elephant shark SIRT3.2 could be influenced by the environmental conditions, we next evaluated the optimal temperature of ΔNT-SIRT3.2 compared to that of human SIRT3. As expected, the optimal temperature of the human SIRT3 was ~37°C (Fig. 9b). In contrast, ΔNT-SIRT3.2 exhibited an optimal temperature at ~24°C (Fig. 9b). The optimal temperature value for the SIRT3.2 deacetylase activity of the elephant shark could represent an evolutionary adjustment to the temperature regime of the habitat in which this species lives, as it has been demonstrated for other enzymes in other species (McCormick 1993; Somero 2004; Bilyk et al. 2021; Saravia et al. 2021). Thus, all subsequent enzymatic activity analyses for ΔNT-SIRT3.2 were performed at 24°C. The apparent specific activity of ΔNT-SIRT3.2 was 260.7 ± 9.9 fluorescence units (FU) min^−1^ mg^−1^, which was significantly lower than that of SIRT3 that was 1143.3 ± 46.3 FU min^−1^ mg^−1^. Apparent *K*_m_ of ΔNT-SIRT3.2 and SIRT3 were 5.7 and 5.5, respectively, which resulted not significantly different (Fig. 9c, d). In contrast, apparent *V*_max_ of ΔNT-SIRT3.2 was 0.322 FU sec^−1^, which was significantly different to that of SIRT3 that was 3.908 FU sec^−1^ (Fig. 9c, d). We next compared the effect of the sirtuin inhibitors nicotinamide (NAM) (Sauve and Schramm 2003) and quercetin (QUE) (You et al. 2019) on the activities of SIRT3 and ΔNT-SIRT3.2. As expected, SIRT3 activity was significantly inhibited by NAM and QUE (Fig. 9e). We also found a significant inhibitory effect of both NAM and QUE on the activity of ΔNT-SIRT3.2 (Fig. 9f). We also compared the effect of the polyphenolic antioxidant resveratrol (REV), which is an activator of sirtuins (Wood et al. 2004). As expected, we found a significant concentration-dependent activation of SIRT3 deacetylase activity up to a 1.6 ± 0.1-fold increase in the presence of 100 μM REV (Fig. 9e). Strikingly, we found a much more potent concentration dependent activation effect of REV on ΔNT-SIRT3.2, showing up to a 11.1 ± 0.6 fold increase in the presence of 100 μM REV (Fig. 9f). However, because the specific activity of ΔNT-SIRT3.2 was ~23% that of SIRT3 in the assay conditions, the activation of ΔNT-SIRT3.2 by REV in fact represents a value only ~1.6 fold higher compared to the activation of SIRT3 by REV. Because different sirtuins have activity on different proteins, as well as remove with different specificity a variety of acyl groups other than the acetyl (Fig. 1) (Anderson et al. 2014), the modulation of ΔNT-SIRT3.2 activity by REV suggests that it has distinct functional substrates. Together, these results demonstrate that elephant shark ΔNT-SIRT3.2 has deacetylase activity and indicate that the activity of SIRT3.2 proteins is modulated in a similar manner to that of mammalian sirtuins.

**Figure 9.**
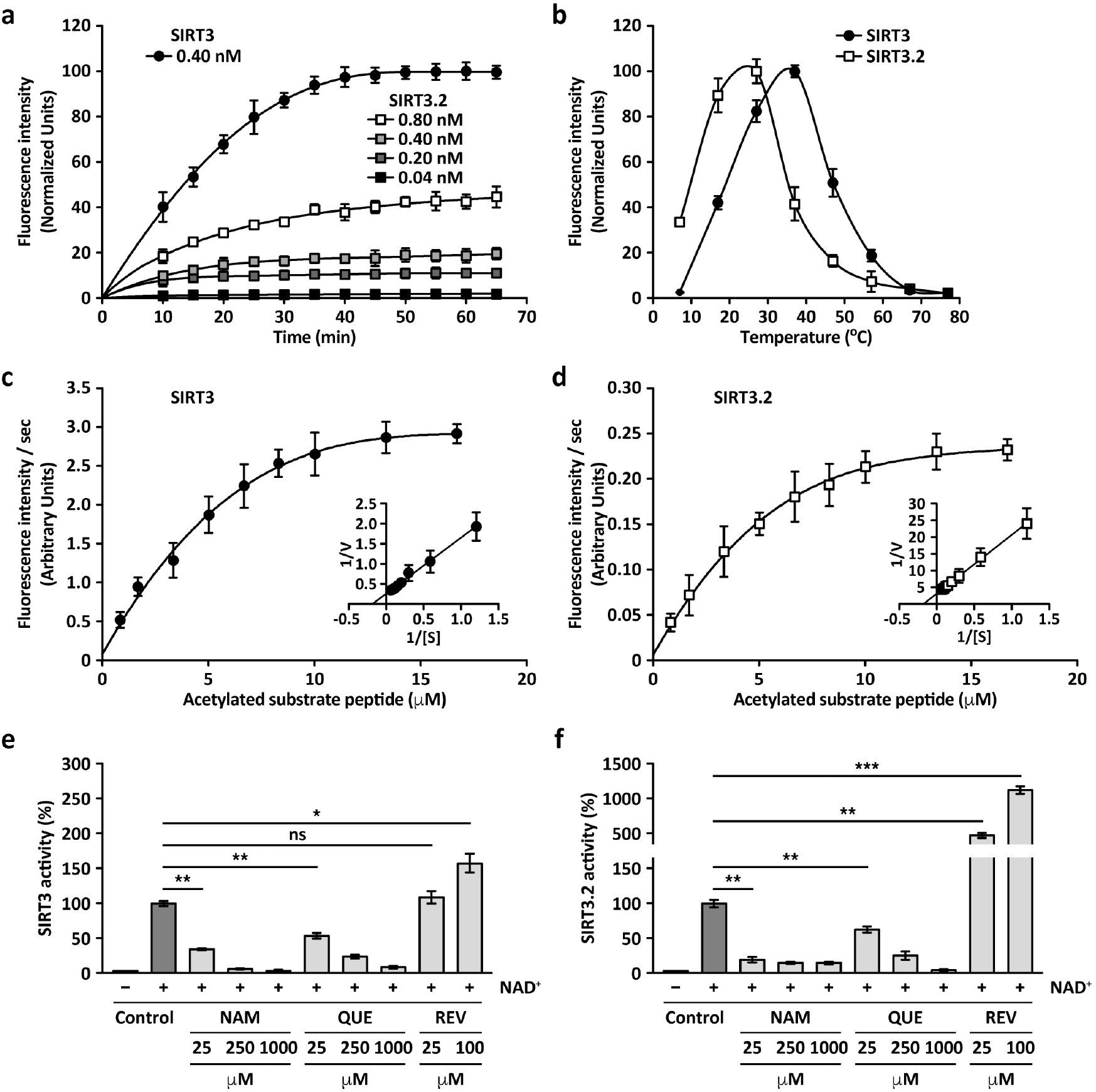
Comparison of deacetylase activity between human SIRT3 and elephant shark ΔNT-SIRT3.2. a-f) Results obtained with a fluorometric deacetylation assay of an acetylated substrate peptide. a) Time course of deacetylase activity at the indicated concentration of SIRT3 and concentrations of ΔNT-SIRT3.2. b) Temperature dependence of deacetylase activity. c-d) Michaelis-Menten plot and Lineweaver-Burk plot of SIRT3 (c) and ΔNT-SIRT3.2 (d) deacetylase activity. e-f) NAD+ dependence and concentration dependent effects of the SIRT3 inhibitors nicotinamide (NAM) and quercetin (QUE) and of the SIRT3 activator resveratrol (REV) on the deacetylase activity of SIRT3 (e) and ΔNT-SIRT3.2 (f). In a-d, graphs depict the mean ± standard deviation (n = 3). In e-f, bars represent the mean ± standard deviation. Statistical analysis was performed using two-tailed unpaired Student’s t-test (n = 3; * p < 0.05; ** p < 0.01; *** p < 0.001; ns, not statistically significant).

#### Elephant shark SIRT3.2 protein increases ATP cellular content

Considering that SIRT3 plays an important role in maintaining basal ATP levels by deacetylation of mitochondrial proteins involved in electron transport chain (ETC), the tricarboxylic acid cycle, and fatty-acid oxidation (Hirschey et al. 2010), we investigated the impact of transient SIRT3.2-3myc expression on ATP cellular content in H4 and HeLa cells (Fig. 10a). Similar to the well-known effect of mammalian SIRT3, SIRT3.2-3myc expression increased ATP cellular content, suggesting that it shares with SIRT3 the ability to enhance the activity of the mitochondrial ETC. Because an enhancement in the activity of ETC can cause an increase in reactive oxygen species (ROS), we also evaluated the levels of ROS in cells expressing SIRT3.2-3myc (Fig. 10b). We found similar ROS levels in cells expressing SIRT3.2-3myc and untransfected cells (Fig. 10b). This finding suggests that SIRT3.2 proteins share the mammalian SIRT3 characteristic of protection against oxidative damage (Bause and Haigis 2013), likely by deacetylation of enzymes involved in ROS clearance to protect mitochondrial membranes from oxidative stress (Tseng et al. 2013). A candidate that could be activated by the deacetylation activity of SIRT3.2 proteins is SOD2 that in human cells reduces cellular ROS content promoting oxidative stress resistance (Qiu et al. 2010; Tao et al. 2010). Alternatively, SIRT3.2 proteins could activate the last complex of the ETC by deacetylation, enhancing ATP-synthase activity without affecting ROS production, similar to the effect that SIRT3 has in response to exercise-induced stress (Vassilopoulos et al. 2014). Together, our results show that the SIRT3.2 protein is a new sirtuin family member with deacetylase activity that could have important roles in the homeostasis of oxidative species and mitochondrial metabolism.

**Figure 10.**
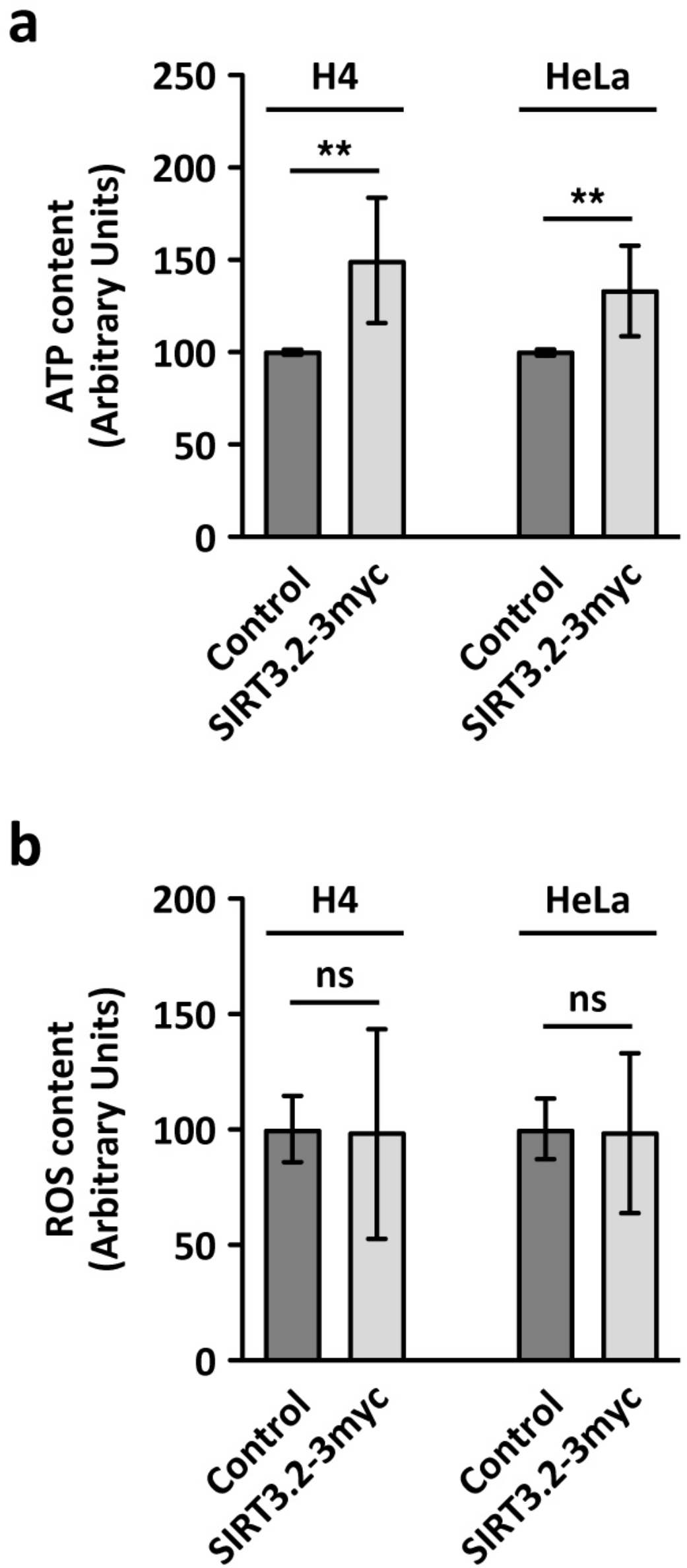
SIRT3.2-3myc expression causes an increment on ATP cellular content with no difference in ROS levels. (a) ATP levels were measured by a luciferin/luciferase bioluminescence assay in H4 and HeLa cells either left untreated (Control) or transiently expressing SIRT3.2-3myc. (b) ROS content was measured by the intensity fluorescence of the dye CM-H2DCFDA in H4 and HeLa cells either left untreated (Control) or transiently expressing SIRT3.2-3myc. Bars represent the mean ± standard deviation. Statistical analysis was performed using two-tailed unpaired Student’s t-test (n = 4; ** p < 0.01; ns, not statistically significant).

#### Conclusions

We performed an evolutionary analysis of the sirtuin gene family in vertebrates. In addition to describing the duplicative history of sirtuins, our phylogenetic analyses revealed the presence of a new gene family member, *SIRT3.2*. This new vertebrate gene lineage is present in all vertebrates other than mammals, birds, and reptiles. Furthermore, our transcriptomic analyses show that *SIRT3.2* is transcribed in almost all examined tissues, showing a tendency to be more expressed in the ovary, suggesting biological functions related to reproduction. According to our analyses, SIRT3.2 localized in the mitochondria, had deacetylase activity and increased the ATP cellular content. The finding of a new sirtuin gene lineage highlights the need for more detailed assessments of orthology with a broader taxonomic sampling to better define the membership composition of gene families (Glover et al. 2019). In the literature, there are examples in which more comprehensive analyses provide a better description of gene families, including previously unknown family members (Castro et al. 2012; Wichmann et al. 2016; Céspedes et al. 2017; Ramos-Vicente et al. 2018; Opazo, Kuraku, et al. 2019; Opazo, Hoffmann, et al. 2019). The availability of species with different gene repertoires could be seen as a natural experiment (Albertson et al. 2009) that helps us understand the evolutionary fate of duplicated genes, as they can fulfill the biological functions associated with the gene family but with a different combination of paralogs.

## Methods

### Protein sequences and phylogenetic analyses

We retrieved sirtuin amino acid sequences in representative species of all main groups of vertebrates. Our sampling included mammals, birds, reptiles, amphibians, lungfish, coelacanth, bony fish, cartilaginous fish, and cyclostomes (Supplementary Table S1). Sequences were obtained from the Orthologous MAtrix project (OMA), December 2021 release (Altenhoff et al. 2021)last accessed on April 8th, 2022). In addition, we complemented our sampling by retrieving sequences from vertebrate groups that were not present or poorly represented in the Orthologous MAtrix project (e.g., crocodiles) from the National Center for Biotechnology Information (NCBI) (Sharma et al. 2019) using the program blast (tblastn) with default parameters (Altschul et al. 1990) Supplementary Table S1). Protein sequences were aligned using the software MAFFT v.7 (Katoh and Standley 2013), allowing the program to choose the alignment strategy (FFT-NS-i, alignment length 2490 amino acids). To select the best-fitting model of molecular evolution we used the proposed model tool in the program IQ-Tree v1.6.12 (Kalyaanamoorthy et al. 2017), which selected JTT+F+I+G4. We used a maximum-likelihood approach to obtain the best tree using the program IQ-Tree v1.6.12 (Trifinopoulos et al. 2016). We carried out 20 independent runs to explore the tree space changing the value of the strength of the perturbation (-pers) parameter. We conducted the following analyses:

1. Five runs modifying the strength of the perturbation parameter from 0.5 (default value) to 0.3.
2. Five runs using the default value (0.5) for the strength of the perturbation parameter.
3. Five runs modifying the strength of the perturbation parameter from 0.5 (default value) to 0.7.
4. Five runs modifying the strength of the perturbation parameter from 0.5 (default value) to 0.9.

In all cases, the number of unsuccessful iterations to stop parameter (-nstop) value was changed from 100 (default value) to 500. The tree with the highest likelihood score was chosen (log-likelihood: −127052.963, -pers 0.5 and -nstop 500). Support for the nodes was evaluated using two approaches: the aBayes test (Anisimova et al. 2011) and the ultrafast bootstrap procedure using 1000 replicates (Hoang et al. 2018). Nicotinamide Nucleotide Transhydrogenase (NNT) protein sequences, a member of the DHS-like NAD/FAD-binding domain superfamily, from the human (*Homo sapiens*), mouse (*Mus musculus*), zebrafish (*Danio rerio*) and spotted gar (*Lepisosteus oculatus*) were used as outgroups.

### Assessment of conserved synteny

We examined genes found upstream and downstream of the sirtuin genes. For comparative purposes, we used the estimates of orthology and paralogy derived from the Ensembl Compara database (Herrero et al. 2016); these estimates are obtained from a pipeline that considers both synteny and phylogeny to generate orthology mappings. These predictions were visualized using the program Genomicus v100.01 (Nguyen et al. 2018). Our assessments were performed in humans (*Homo sapiens*), chicken (*Gallus gallus*), gharial (*Gavialis gangeticus*), red-eared slider (*Trachemys scripta*), green anole (*Anolis carolinensis*), tropical clawed frog (*Xenopus tropicalis*), coelacanth (*Latimeria chalumnae*), spotted gar (*Lepisosteus oculatus*), and elephant shark (*Callorhinchus milii*).

### Dot-plots

We retrieved the chromosomal region containing the *SIRT3.2* gene of the tropical clawed frog (*Xenopus tropicalis*, Ch3) including 6.7 kb up- and downstream, and the corresponding syntenic region in the human (*Homo sapiens*, Chr12), opossum (*Monodelphis domestica*, Chr8), chicken (*Gallus gallus*, Chr1), gharial (*Gavialis gangeticus*, NW_017729022.1), red-eared slider (*Trachemys scripta*, Chr1), and green anole (*Anolis carolinensis*, Chr5) based on the location of the flanking genes (*RIC8B* and *TMEM263*). We aligned SIRT3.2 syntenic regions using Advanced PipMaker (Schwartz et al. 2000).

### Transcript abundance analyses

Sirtuin transcript abundance was measured from a representative sample of vertebrates including the elephant shark (*Callorhinchus milii*), zebrafish (*Danio rerio*), tropical clawed frog (*Xenopus tropicalis*), anole lizard (*Anolis carolinensis*), and human (*Homo sapiens*). RNASeq libraries from brain, heart, kidney, liver, muscle, ovary, and testis from each species were gathered from the NCBI Short Read Archive (SRA; (Leinonen et al. 2011). Accession numbers for species and tissue specific libraries can be found in Supplemental Table S2. Reference transcript sequences were collected from Ensembl v.100 (Yates et al. 2020) and we removed sequences that were shorter than 100 bp. For each library adapters were removed using Trimmomatic 0.38 (Bolger et al. 2014) and reads were filtered for quality using the parameters HEADCROP:5, SLIDINGWINDOW:5:30, and MINLEN:50. We mapped quality filtered paired-end RNAseq reads back to reference sequences using Bowtie 1.2.2 (Langmead et al. 2009) and default parameters of RSEM (Li and Dewey 2011). Transcripts with < 10 mapped reads across all seven tissues per species were removed prior to normalization. Normalization of raw read counts for each species was performed using the median of ratios (Anders and Huber 2010) method implemented by the estimateSizeFactors function in DESeq2 v1.26 (Love et al. 2014). Briefly, for all samples within a species, the geometric mean is calculated for the read counts of each gene. Read counts are then divided by the geometric mean and the median ratio is determined for each sample. Normalized read counts are calculated by dividing raw read counts by the sample-specific median ratio. If multiple SIRT transcripts were present, we presented the expression data from the transcript with the most mapped reads.

### Protein structure homology modeling

Protein structure homology modeling was performed using the SWISS-MODEL server (https://swissmodel.expasy.org/) (Waterhouse et al. 2018). Structural figures were prepared with PyMOL Molecular Graphics System, Version 2.0.6 Schrödinger, LLC.

### Recombinant cDNA Constructs

For expression in mammalian cells, a codon-optimized mammalian expression construct encoding full-length (amino acids 1-372) elephant shark (*Callorhinchus milii*) SIRT3.2 (Supplementary table S1), cloned in-frame into the *Eco*RI and *Sal*I sites of the pCMV-3Tag-4a vector followed by vector encoding sequence of three successive Myc epitope tags before a stop codon, was acquired from GenScript (Piscataway, NJ). For the generation of a codon-optimized bacterial expression construct, a cDNA encoding N-terminal truncated (amino acids 95-372) elephant shark SIRT3.2, cloned in-frame into the *Eco*RI and *Sal*I sites of the pGEX-4T-1 vector, was also acquired from GenScript. This cDNA was subsequently cloned in-frame into the *Eco*RI and *Sal*I sites of the pGST-Parallel-1 vector (Sheffield et al. 1999). The nucleotide sequence of all recombinant constructs was confirmed by dideoxy sequencing using the AUSTRAL-omics core facility at Universidad Austral de Chile (https://australomics.cl/).

### Cell culture, cell transfection and preparation of protein extracts

H4 human neuroglioma cells, HeLa human cervix adenocarcinoma cells and MDA-MB-231 human mammary gland adenocarcinoma cells were obtained from the American Type Culture Collection (Manassas, VA). H4 and HeLa cells were maintained in Dulbecco’s modified Eagle’s medium (DMEM), and MDA-MB-231 in DMEM-F12 (ThermoFisher, Waltham, MA). For all cell lines, media were supplemented with 10% heat-inactivated fetal bovine serum, 100 U/ml penicillin, 100 μg/ml streptomycin (ThermoFisher), and 5 μg/ml plasmocin (InvivoGen, San Diego, CA), and cells were cultured in a humidified incubator with 5% CO_2_ at 37 °C. For transient transfections, cells were either seeded on top of glass coverslips on 24-well plates or on 6-well plates. When cells were ~60% confluent, transfections were performed with Lipofectamine 2000 (ThermoFisher), according to the manufacturer’s instructions. Preparation of protein extracts from cultured, transfected or non-transfected cells was performed using methods described elsewhere (Tenorio et al. 2016).

### Antibodies and cell reagents

We used the following antibodies: mouse clone 9E10 against the c-Myc epitope (Covance, Princeton, NJ), mouse clone BA3R against β-ACTIN (ThermoFisher), rabbit polyclonal against Cytoskeleton-associated protein 4 (P63; Merck, Germany; cat # HPA001225), and sheep polyclonal against Trans-Golgi network integral membrane protein 2 (TGN46; Bio-Rad Laboratories, Hercules, CA; cat # AHP500G). The following fluorochrome-conjugated antibodies were from ThermoFisher: Alexa Fluor-488– conjugated donkey anti mouse IgG, Alexa Fluor-594–conjugated donkey anti rabbit IgG and Alexa Fluor-647-conjugated donkey anti sheep IgG. HRP-conjugated secondary antibodies were from Jackson ImmunoResearch (West Grove, PA). Depending on their reactivity, primary antibodies were used at a dilution 1/200 to 1/2000. HRP-conjugated secondary antibodies were used at dilutions 1/5000 to 1/20000 also depending on their reactivity. All Alexa Fluor-conjugated secondary antibodies were used at a dilution 1/1000. The fluorescent nuclear stain 4’,6-diamidino-2-phenylindole (DAPI) was also from ThermoFisher.

### Protein Electrophoresis, Immunoblotting and Immunofluorescence Microscopy

SDS-PAGE and immunoblotting were performed as described (Bustamante et al. 2020). For fluorescence microscopy, cells grown on glass coverslips were transfected, and after 16-h cells were left untreated or treated for 30 min with MitoTracker™ Orange CMTMRos (ThermoFisher) according to the manufacturer’s instructions. After washing with phosphate buffered saline (PBS) supplemented with 0.1 mM CaCl2 and 1 mM MgCl2 (PBS-CM), cells were fixed in 4% paraformaldehyde for 30 min at room temperature, permeabilized with 0.2% Triton X-100 in PBS-CM for 15 min at room temperature, and incubated with blocking solution (0.2% gelatin in PBS-CM) for 10 min at room temperature. Cells were incubated simultaneously either with antibodies to the c-Myc epitope (1/200) and to TGN46 (1/1000) or to the c-Myc epitope, to TGN46 and to P63 (1/600) for 30 min at 37 °C in a humidified chamber. After washing coverslips with PBS, cells were incubated as before, but with fluorochrome-conjugated antibodies, followed by PBS-washing, and coverslips were mounted onto glass slides with Fluoromount-G mounting medium (ThermoFisher). Fluorescence microscopy images were acquired with an AxioObserver.D1 microscope equipped with a PlanApo 63x oil immersion objective (NA 1.4), and an AxioCam MRm digital camera (Carl Zeiss). To quantitatively evaluate colocalization of fluorescence signals, we obtained the Pearson’s correlation coefficient and cytofluorograms of pairwise comparisons from images acquired under identical settings, avoiding signal saturation and corrected for background, crosstalk and noise signals on each set of images, using the plugin JACoP (Bolte and Cordelières 2006) implemented in the software ImageJ (version 1.47h)(Schneider et al. 2012), and the plugin Colocalization Finder implemented in the software ImageJ2 (version 2.3.0/1.53) (Rueden et al. 2017). To prepare figures, 12-bit images were processed with ImageJ software and Adobe Photoshop CS3 software (Adobe Systems, Mountain View, CA).

### Expression and Purification of Recombinant N-terminal truncated elephant shark SIRT3.2 protein

Recombinant, N-terminal truncated elephant shark SIRT3.2 (ΔNT-SIRT3.2; amino acids 95-372) tagged with an N-terminal glutathione S-transferase (GST) followed by a tobacco etch virus (TEV) protease cleavage site was expressed and purified using a method described previously (Ross et al. 2014), with some modifications. Briefly, expression in *E. coli* B834(DE3) (Novagen, Madison, WI) was induced with 0.25 mM IPTG at 25 °C for 16 h. Pellets of bacteria were resuspended in homogenization buffer (50 mM Tris HCl, 0.5 M NaCl, 5 mM EDTA, 5 mM β-mercaptoethanol, and 2 mM phenylmethylsulfonyl fluoride, pH 8.0) and lysed by sonication. The clarified supernatant was purified on glutathione-Sepharose 4B (GE Healthcare). After removal of the GST moiety by TEV cleavage, and sequential further passage through glutathione-Sepharose 4B and Ni-NTA (QIAGEN) resins, ΔNT-SIRT3.2 was further purified on a Superdex 200 pg column (GE Healthcare) equilibrated in storage buffer (50 mM Tris HCl, 150 mM NaCl, 1 mM DTT). Aliquots of purified protein were kept at −80 °C until use.

### Deacetylation activity

To characterize deacetylation activity of recombinant, N-terminal truncated elephant shark SIRT3.2 (ΔNT-SIRT3.2), we used a fluorometric SIRT3 activity assay kit (Abcam, cat # ab156067; Cambridge, UK), according to the manufacturer’s instructions. This assay allows detection of a fluorescent signal upon deacetylation of an acetylated substrate peptide for recombinant human SIRT3. The intensity of fluorescence was measured on a fluorometric microplate reader (Varioskan Flash, ThermoFisher) with excitation set at 350 nm and emission detection set at 450 nm. To analyze the deacetylation activity of N-terminal truncated elephant shark SIRT3.2 protein, first we determined its optimum temperature that resulted to be ~24 °C. Subsequent analyses were performed by comparing the activity of N-terminal truncated elephant shark SIRT3.2 protein at 24 °C to that of human SIRT3, which is provided by the assay kit, at 37 °C. *K*_m_ and *V*_max_ were obtained by varying the concentration of the fluorogenic acetylated substrate peptide provided by the assay kit. This assay was also used to determine the effect of nicotinamide (NAM), quercetin (QUE) and resveratrol (REV; Sigma-Aldrich) on the deacetylation activity of ΔNT-SIRT3.2.

### ATP and ROS content

ATP levels were quantified in cell lysates using a luciferin/luciferase bioluminescence assay (ATP determination kit, Molecular Probes cat # A22066, Thermo Fisher Scientific) as previously described (Jara et al. 2018). Briefly, cells were lysed in HEPES buffer (125 mM NaCl, 25 mM NaF, 1 mM EDTA, 1 mM EGTA, 1% NP-40, 25 mM HEPES, pH 7.4) supplemented with a protease inhibitor mixture (cat # 78429, Thermo Fisher Scientific) and centrifuged at 12000 rpm using a refrigerated centrifuge (Eppendorf model 5424R) for 15 min at 4 °C. Proteins in the supernatant were quantified, and 10 μl of lysate were used to determine ATP levels according to the kit manufacturer’s instructions. The ATP level in each sample was calculated using standard curves and normalized to the protein concentration in each sample. ROS content was determined in cell lysates using 25 μM CM-H2DCFDA (DCF; a fluorogenic indicator of ROS), as previously described (Jara et al. 2018; Olesen et al. 2020). The intensity of fluorescence was measured on a fluorescence plate reader (Biotek Synergy HT, Agilent, Santa Clara, CA) with excitation set at 485 nm and emission detection set at 530 nm. Briefly, in a dark 96-well plate, 25 μg of cell lysate proteins were incubated with CM-H2DCFDA (DCF) dye with shaking for 5 min at room temperature, followed by determination of the fluorescence of each sample that was subtracted by the fluorescence of the respective blank.

### Statistical Analysis

In the case of the deacetylation, ATP content and ROS content assays, quantification was performed from at least three independent experiments/measurements. Statistical analysis was performed using Microsoft Excel for Mac 2011 (Microsoft Corporation). When appropriate, results are represented in graphs depicting the mean ± standard deviation. Statistical significance was determined by two-tailed, paired t-test. P-values > 0.05 or ≤ 0.05 were regarded as not statistically significant or statistically significant, respectively. In the figures, P-values between 0.01 and 0.05 are indicated with one asterisk, P-values between 0.001 and 0.01 are indicated with two asterisks, and P-values less than 0.001 are indicated with three asterisks.

## Acknowledgements

This work was supported by the Fondo Nacional de Desarrollo Científico y Tecnológico from Chile (FONDECYT 1210471) and The Millennium Nucleus of Ion Channel-Associated Diseases is a Millennium Nucleus of the Millennium Scientific Initiative, National Agency of Research and Development (ANID, NCN19_168), Ministry of Science, Technology, Knowledge and Innovation, Chile to JCO, Fondo Nacional de Desarrollo Científico y Tecnológico from Chile (FONDECYT 1211481) to GAM, National Science Foundation (EPS-0903787, DBI-1262901 and DEB-1354147) to F.G.H., Financiamiento Basal No. ANID/BASAL/ACE210009 to PVB and No. ANID/BASAL/FB210008 to PVB and CTR, Fondap-Ideal Grant N° 15150003 and ANID-Millennium Science Initiative Program-Center ICM-ANID ICN2021_002 to LV-C, Fondo Nacional de Desarrollo Científico y Tecnológico from Chile (FONDECYT 1180957) to FJM, Vicerrectoría de Investigación, Desarrollo y Creación Artística, Universidad Austral de Chile to JCO, FJM, LVC, GAM. We thank Stefanie Teuber, John Quiroga and Rafael Burgos, Instituto de Farmacología y Morfofisiología, Facultad de Ciencias Veterinarias, Universidad Austral de Chile, for providing assistance in the use of the Varioskan Flash microplate reader; Hianara Bustamante, Instituto de Microbiología Clínica, Facultad de Medicina, Universidad Austral de Chile, for providing quercetin and resveratrol; Víctor Castro, Departamento de Biología, Facultad de Ciencias, Universidad de Chile, for providing NAD^+^ and nicotinamide; and Gonzalo Astroza, and Andrés Rivera-Dictter for technical assistance.

## Contributions

JCO and GAM designed the study. JCO, MWV, FGH, KZ, CM, CL, VC, FJM, PVB, CTR, GAM, collected and/or analyzed data. JCO, PVB and GAM wrote the manuscript. JCO, MWV, FGH, KZ, CM, CL, VC, LV-C, FJM, PVB, CTR and GAM, reviewed and edited the manuscript. All authors contributed to the article and approved the submitted version.

## Ethics declarations

### Data availability

Data and supplementary material are available online at zenodo.org

